# Platinized graphene fiber electrodes revealed differential activity in terminal splenic neurovascular plexi supporting a direct spleen-vagus communication

**DOI:** 10.1101/2021.03.19.436189

**Authors:** María A. González-González, Geetanjali Bendale, K Wang, Gordon G. Wallace, Mario Romero-Ortega

## Abstract

Neural interfacing nerve fascicles along the splenic neurovascular plexi (SNVP) is needed to better understand the spleen physiology, and for selective neuromodulation of this major organ. However, the small size and anatomical location have proven to be a significant challenge. Here, we use a reduced liquid crystalline graphene oxide (rGO) fiber coated with platinum (Pt) as a super-flexible suture-like electrode to interface multiple SNVP. The Pt-rGO fibers were used as handover knot electrodes over the small plexi, allowing sensitive recording from the splenic nerve (SN) terminal branches. Asymmetric defasciculation of the SN branches was revealed by electron microscopy, and the functional compartmentalization in spleen innervation was evidenced in response to hypoxia and pharmacological modulation of mean arterial pressure. We demonstrate that electrical stimulation of cervical and subdiaphragmatic vagus nerve (VN), evoked direct activity in a subset of SN terminal branches, providing evidence for a direct VN control over the spleen. This notion was supported by tract-tracing of SN branches, revealing an unconventional direct spleen-VN projection. High-performance Pt-rGO fiber electrodes, may be used for the fine neural modulation of other small neurovascular plexi at the point of entry of the organs as a bioelectronic medical alternative.

## Introduction

Vagus nerve stimulation (VNS) has been shown to reduce the circulation of inflammatory cytokines and to improve survival during severe inflammation in rodents ^1-3^. The immunosuppression effect of VNS seems to be mediated by activation of the inflammatory reflex pathway, where efferent VN innervating the celiac ganglia (CG) form the splenic nerve (SN), predominantly by sympathetic vasoconstrictor fibers ^4,5^, and mediate the release of acetylcholine (Ach) from T-cells in the spleen ^6,7^. These studies led to the notion that VNS can be used as a treatment for severe immune disorders, including inflammatory bowel diseases and rheumatoid arthritis ^2,8^. However, the idea of cholinergic VN-CG-SN innervation to the spleen is a subject of much debate, and was directly challenged by Bratton and collaborators who failed to confirm a direct SN synaptic activation by VNS ^9^. The controversy is also based on the apparent lack of sensory input to the spleen, and the consideration that its neural control is exclusively sympathetic ^10^. It is known that efferent norepinephrine (NE) fibers travel along arterial and apical branches to innervate lymphocytes and macrophages in the spleen ^11,12^. However, a small percentage of Ach fibers are located apically ^13^, and seem to represent afferent axons from vein baroreceptors projecting to the nucleus of the tractus solitarious in the brain stem, which responds to changes in blood pressure (BP) ^14-16^. Indeed, the SN contains axons from different peptidergic neurons that were proposed to differentially modulate spleen vascular functions ^17,18^. Part of the challenge is the small size of the rat SN (i.e., ∼100μm diameter) composed of 300-400 axons in 3-6 fascicles traveling along blood vessels, and forming a neurovascular plexi (SNVP), that branches before entering the 2-3 lobular organ ^4,19^. This makes the visualization and isolation of the SN for electrophysiology or tract-tracing studies, highly challenging.

We reasoned that the functional understanding of the complex innervation to the spleen will require simultaneous and independent recording of the four splenic terminal neurovascular plexi (SNVP-1 to 4) during evoked physiological events. However, current silver or platinum hook or cuff electrodes lack the miniaturization, flexibility, sensitivity, or charge injection capacity to allow effective communication with these small nerve fascicles ^13,20^. We recently developed a reduced liquid crystalline graphene oxide (rGO) fiber coated with a thin platinum (Pt) layer to produce a highly sensitive and super-flexible Pt-rGO electrode with unmatched mechanical and electrochemical characteristics ^21^. The high geometric surface area of these fiber electrodes provides unrivaled charge injection capacity (10.34 mC/cm^2^) and low impedance (28.4 ± 4.1 MΩ µm^2^; 1kHz), superior to conventional carbon nanotube arrays or other common materials used as neural interfaces ^22,23^, and able to record cortical neural activity with a signal to noise ratio (SNR) of 9.2dB. Here, we report that Pt-rGO fibers can be used as an overhand knot suture electrode (aka “sutrode”) around individual SNVP, without isolating the SN terminal branches from the adjacent vasculature, to effectively and sensitively interface this nerve. Transmission electron microscopy (TEM) of the SN branches revealed asymmetric defasciculation from the parent nerve, and simultaneous neural recording from all terminal SN branches using the sutrode, revealed functional compartmentalization in spleen innervation in response to hypoxia and pharmacological modulation of BP. Furthermore, electrical stimulation of cervical and subdiaphragmatic VN (cVN and SD-VN), evoked direct activity in a subset of SN terminal branches, providing evidence for a direct VN control over the spleen. This notion was supported by tract-tracing of apical and basal SN branches, revealing an unconventional direct spleen-VN projection. The use of high-performance Pt-rGO fiber electrodes provided functional information of individual small SNVP, shedding light on the fine neural control of internal organ physiology and enabling neuromodulation at the point of entry.

## Results

### Use of Pt-rGO fibers as “sutrodes”

The flexibility and mechanical robust nature of the Pt-rGO fibers suggested its use as a suture electrode. To that end, we made a slight modification to the previously reported fabrication method ^21^, masking 1 cm proximal and distal segments of 10 cm Pt-rGO fibers before Parylene C coating. The distal end was soldered to a Pt wire using silver (Ag) paste for connection to amplifiers, and the distal end was used for recording/stimulation (Fig. 1a-a’). Cyclic voltammetry was used to confirm that the electrochemical characteristics of the Pt-rGO fibers were not affected by these changes (Fig. 1b). We then measured the impedance of the sutrode while tying a knot to complete closure and confirmed the flexibility and conductivity of the fiber electrode (Fig. 1c; Supplementary video 1). The noise measured by the sutrode was lower compared to that sensed by a conventional Pt hook electrode (Fig. 1e). To demonstrate the mechanical robustness of the sutrode, we tied it to a 9-0 polyamide suture needle, and successfully drove the sutrode through the rat biceps femoris muscle without breaking (Fig. 1f). As expected, passing an electrical current of 2-10 V potential, evoked visible graded muscle contractions (Supplementary video 2). We then evaluated the use of the sutrode as a recording electrode by wrapping it around the sciatic nerve, tying a knot over it carefully, not to occlude the epineural blood circulation (Fig. g), and separately, used as a monopolar electrode to record the evoked compound action potential from muscle sensory afferents as the muscle contracted were recorded in the ScN (Fig, 1h). We then placed the sutrode around the tibial nerve fascicle and using a stimulating hook electrode proximally, demonstrated the recording of compound nerve action potentials (CNAPs) evoked at different voltage strengths (Fig. 1i-j). In addition, and consistent with our previous report in the cerebral cortex, we also confirmed that multiple individual sutrodes can be inserted directly into peripheral nerves for a sensitive recording of intraneural single units with a SNR= 9.6 (Supplementary Figure 1). These tests confirmed the use of the sutrode as a sensitive overhand knot electrode for modulation of the peripheral nerve.

**Figure 1.**
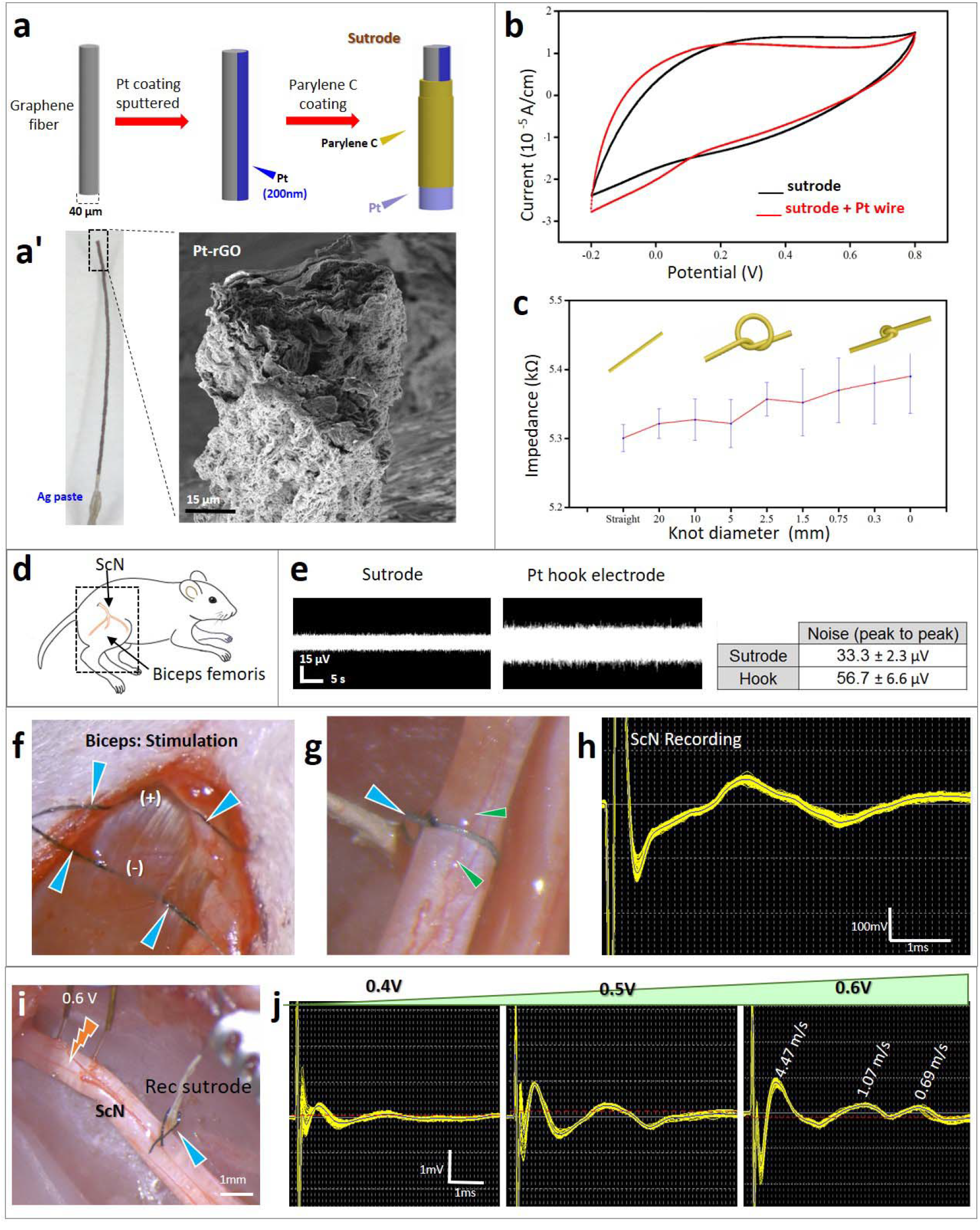
Platinized graphene fibers serve as suture and wrapping electrodes. a) Coating steps of extruded Pt-rGO electrodes. a’) photograph and SEM. b) Cyclic voltammetry of Pt-rGO fibers before and after soldering to a Pt wire. c) Graphene fiber knotting does not alter the electrode impedance. d) Illustration of the ScN and biceps femoris muscle. e) Reduced baseline noise of the sutrode compared to Pt hook electrodes in saline. f) Sutrode placed on biceps (blue arrows) evoked effective muscle contraction (Supplementary Video 1). g) Sutrode wrapped snugly on the ScN (blue arrow), without blood flow occlusion of superficial vasculature (green arrows) used to h) record the evoked CNAP. i) Stimulation of the tibial nerve fascicle with hook electrodes and recording with the sutrode (blue arrow), allowed the recording of j) graded evoked CNAPs and the detection of B and C fibers. Pt-rGO, reduced liquid crystalline graphene oxide, ScN, sciatic nerve, SEM, scanning electron microscopy; CNAP, compound nerve action potentials.

### Sutrode recording of physiologically evoked activity in the cVN

We tested the ability of the sutrode to record spontaneous and evoked physiological activity in the cVN using the Pt-rGO fiber as a hook electrode. Stable baseline activity was recorded with 20 µV peak-to-peak (pp; biological noise) and spontaneous waveforms of 30-60 µV pp at 1-3 spikes/sec identified by principal component analysis (PCA) (Supplementary Fig. 2). We then induced hypotension by the intravenous administration of the nitric oxide donor sodium nitroprusside (NPS) to evaluate the evoked neurophysiological responses in the cVN (Fig 2a). After □1 min of NPS administration, the mean arterial blood pressure (MAP) decreased by 50 mmHg approximately. This coincided temporally with a reduction in the amplitude of spontaneous activity in vagal activity lasting 20 seconds, which was followed by the appearance of compound nerve action potentials (CNAPs) activity of 80-100μV waveforms at high frequency (Fig. 2b), likely from baroreceptor and cardiovascular afferents. To confirm the ability to record physiological activity from different types of axons in the cVN, changes in spontaneous activity were evoked by 2 min lapse in oxygen restriction. Fig. 2c shows a representative recording of basal spontaneous activity of an isolated compound action potential waveform (Fig. 2d). This activity decreased immediately after the hypoxia, but 180 seconds later a new CNAP was detected firing at 10-fold higher frequency compared to baseline, likely the result of cardio- pulmonary reflex activity, as it was reversed by normoxia (Fig. 2c-d). The initial decrease in cVN activity one minute after Oxygen restriction correlated with a mild reduction in heart rate (from 348 to 341±4.1 beats per minute) followed by a mild increase in breathing rate (from 42 to 46±2.8 breaths per minute) (Fig. 2e-f), suggesting the initial reduction in baroreceptor signals and subsequent recruitment of lung Aδ/B mechanoceptors and nociceptors. Together, the data confirmed the use of the sutrode as a sensitive electrode for extraneural recording of physiological compound action potentials in the cVN.

**Figure 2.**
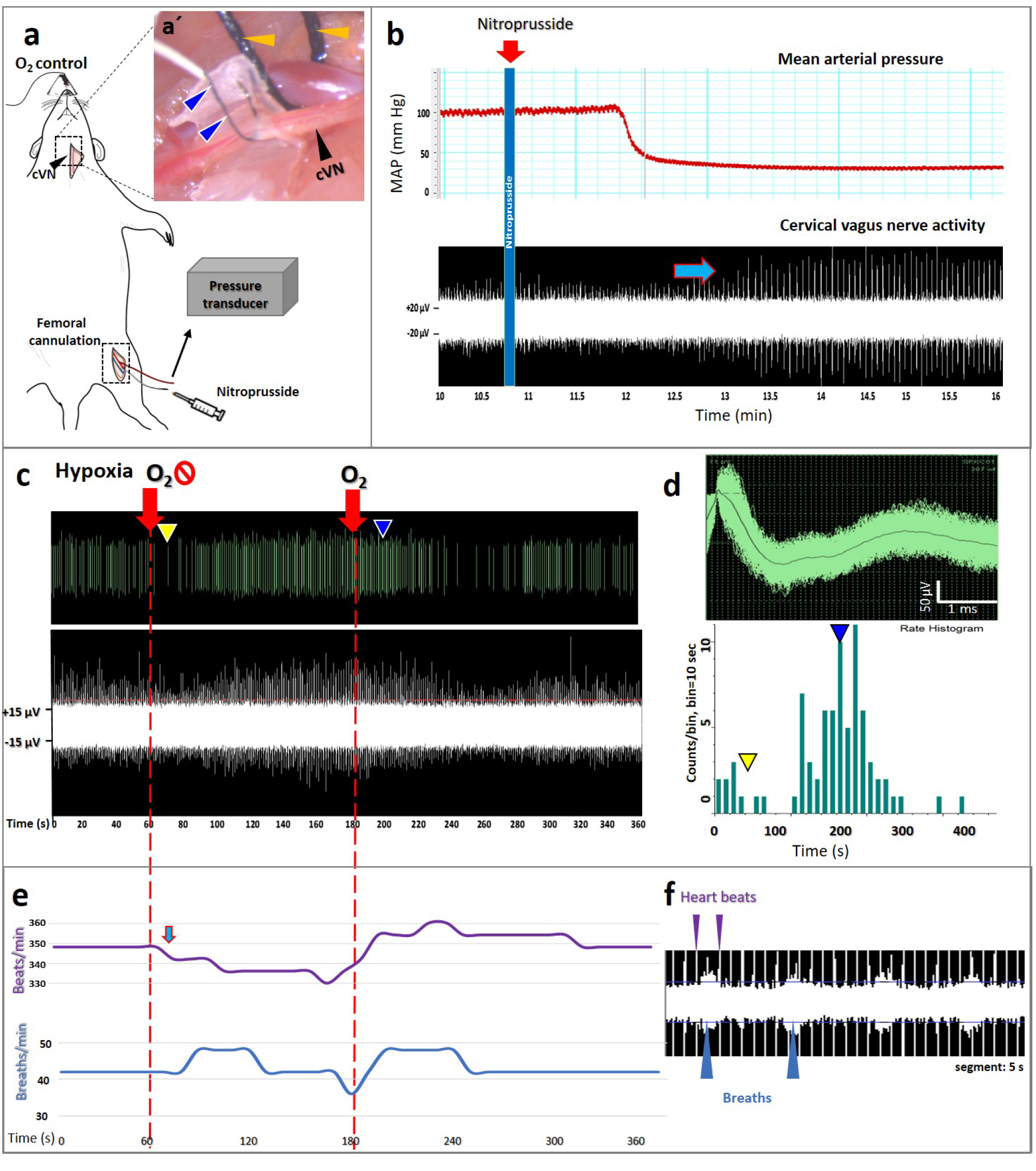
Vagal neuronal activity evoked by hypotension and hypoxia, recorded by sutrode. a) Schematic of sutrode placement on the rat cVN, with the femoral arterial blood pressure sensor. a’) The cVN (black arrow) was interfaced with a sutrode (blue arrows). A conventional 4-0 nylon suture was used to isolate the carotid artery (yellow arrows). b) Nitroprusside (red arrow) induced hypotension, which correlated with changes in cVN firing rate (blue arrow). c) Oxygen restriction induced an initial decrease (yellow arrowheads) and subsequent 10-fold increase (blue arrowheads) in vagal activity confirmed by d) rate histogram of an isolated waveform. Oxygen restriction and restoration are pointed as red arrows in c. e) The initial reduction corresponded to a slight decrease in heart rate (blue arrow), and a subsequent increase in respiration rate calculated from f) unfiltered raw data files. cVN, cervical vagus nerve; MAP, mean arterial pressure.

### Asymmetric axon branching into terminal splenic neurovascular plexi

Functional studies on the neural control of the spleen are limited by the location of the SN, as it travels along the vasculature and splits into terminal branches before entering into the organ (Fig. 3a) and the incomplete anatomical description of the axonal fasciculation. To learn about the axon composition in the SN branches, we did histological and electron microscopic evaluation of the terminal four SNVP, and observed multiple fascicles in the SN ranging from 100-500 μm in diameter, located between the splenic artery and vein, and composed of a heterogeneous population of axon types with the majority being unmyelinated (Fig. 3b-c). The SN fascicles split along the vascular branches into 1-2 fascicles (50-100 μm in diameter) that followed the splenic vasculature and surrounded by fatty tissue (Fig. 3d). Morphometric analysis of the SNVP, revealed that the splitting of SN axons into the terminal branches is notably asymmetric (Fig. 3d-e), where the average diameter of unmyelinated axons was 1.70±0.4 μm in SN-1, 0.92±0.2 μm in SN-2, 1.38±0.4 μm in SN-3, and 1.89±0.4 μm in SN-4. Furthermore, few (3-6) myelinated axons were present only in SN-1 (median 2.54 μm; 0.85 g-ratio) and SN-2 (median 0.95 μm; 0.71 g-ratio), but not in SN-3,4. The different axon composition is evident in Fig. 3f, where a normality test revealed that SN-2 showed normal distribution in axon diameter (p<0.001), whereas others do not (Fig. 3f). Goodness to fit test showed that SN-2 and SN-4 have significantly different slopes (F = 2.87. DFn = 4, p<0.05; Fig. 3f), indicating that the axon composition of the terminal branches is not homogeneous (Fig. 3 g-h).

**Figure 3.**
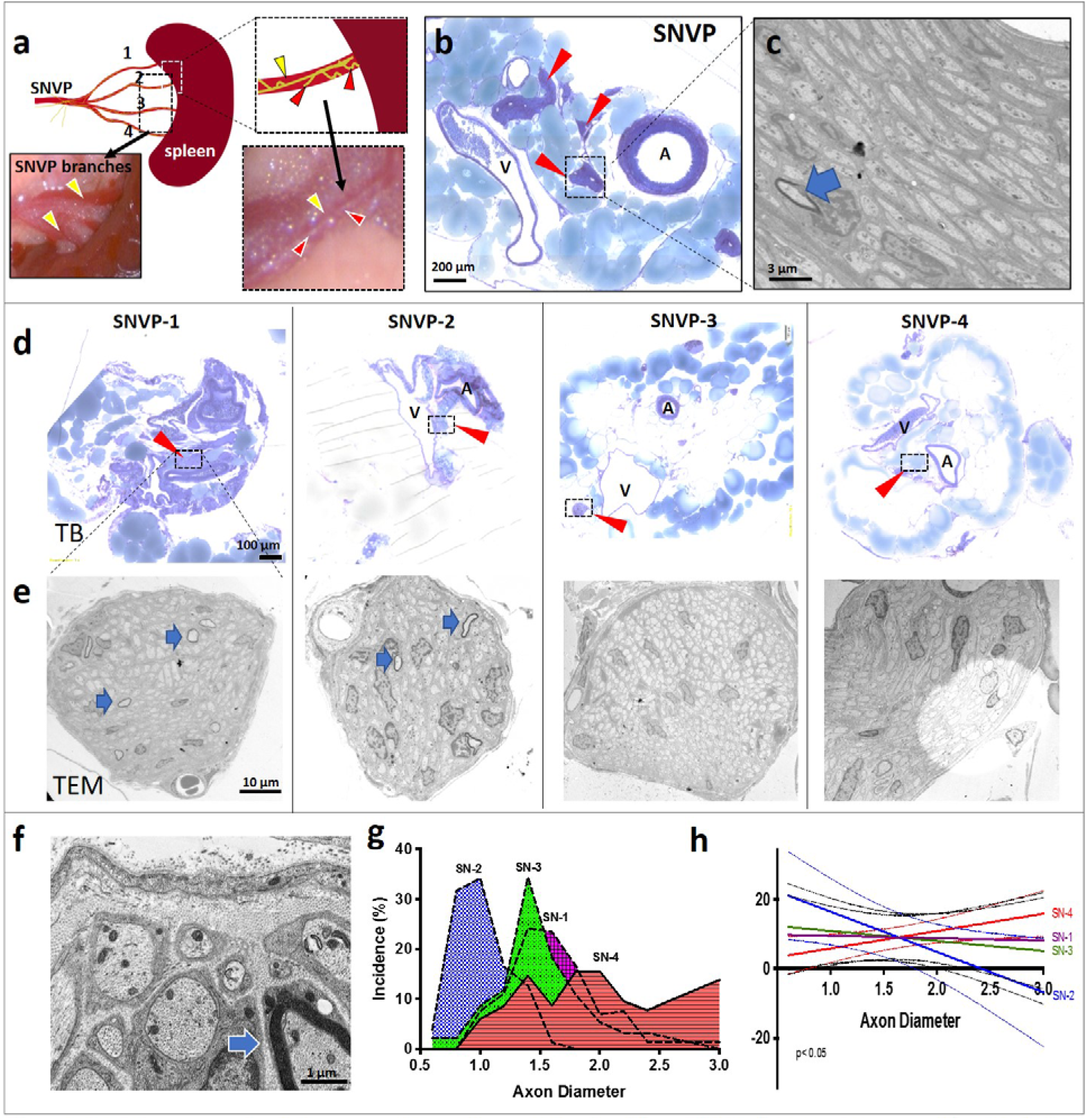
Splenic neuro-vascular plexi asymmetric branching. a) Schematic of the rat SNVP splitting into four terminal branches before entering into the spleen. Inserts show the splenic vasculature (yellow arrowheads) and the SN (red arrowheads) b) Toluidine blue (TB) staining of the SNVP, red arrows point the location of the SN fascicles between the artery (A) and vein (V). c) unmyelinated SN axons by TEM with a single myelinated axon (blue arrow). d) TB staining of the four terminal SN branches, with nerve fascicles in inserts (red arrowheads) e) TEM micrographs of axonal composition in each terminal branch. Myelinated axons (blue arrows) were observed in SN1-2, but not in SN 3-4. f) High TEM magnification shows the detailed SN-2 axons. g) Axon diameter distribution in the SN branches. h) Linear regression analysis of axon diameter distribution. SN, splenic nerve; SNVP, splenic neurovascular plexi, TEM, transmission electron microscopy. The slope between SN-1 and SN-4 is significantly different (p<0.05).

### Sutrode interfacing of splenic terminal neurovascular plexi revealed differential physiology

The heterogeneous axonal composition in the splenic terminal neurovascular plexi (SNVP) compelled us to use the sutrode to record simultaneously from the four terminal branches and investigate if their neural activity response varied in response to specific physiological stimuli. To that end, we made a handover knot with the sutrode to wrap and interface each terminal splenic branch (Fig. 4a, a’). Despite the small size (50-100 μm) and inter-vessel location of these nerve fascicles, we were able to record for the first time, spontaneous activity with great sensitivity (SNR= 8.5±0.2 pp, Fig. 4b and b’). We used PCA on the unfiltered data to identify neural waveforms of spontaneous activity in a 6.5 ms window. Raster plot of the selected units (Fig. 4c-d) showed distinct waveforms ranging from 50 μV pp in SN-1,3 to 400 μV pp in SN-4 (Fig. 4d). The pattern of spontaneous activity in the four SN terminal branches seems different, with tonic activity observed in SN-1 and SN-3, but not in the other two branches.

**Figure 4.**
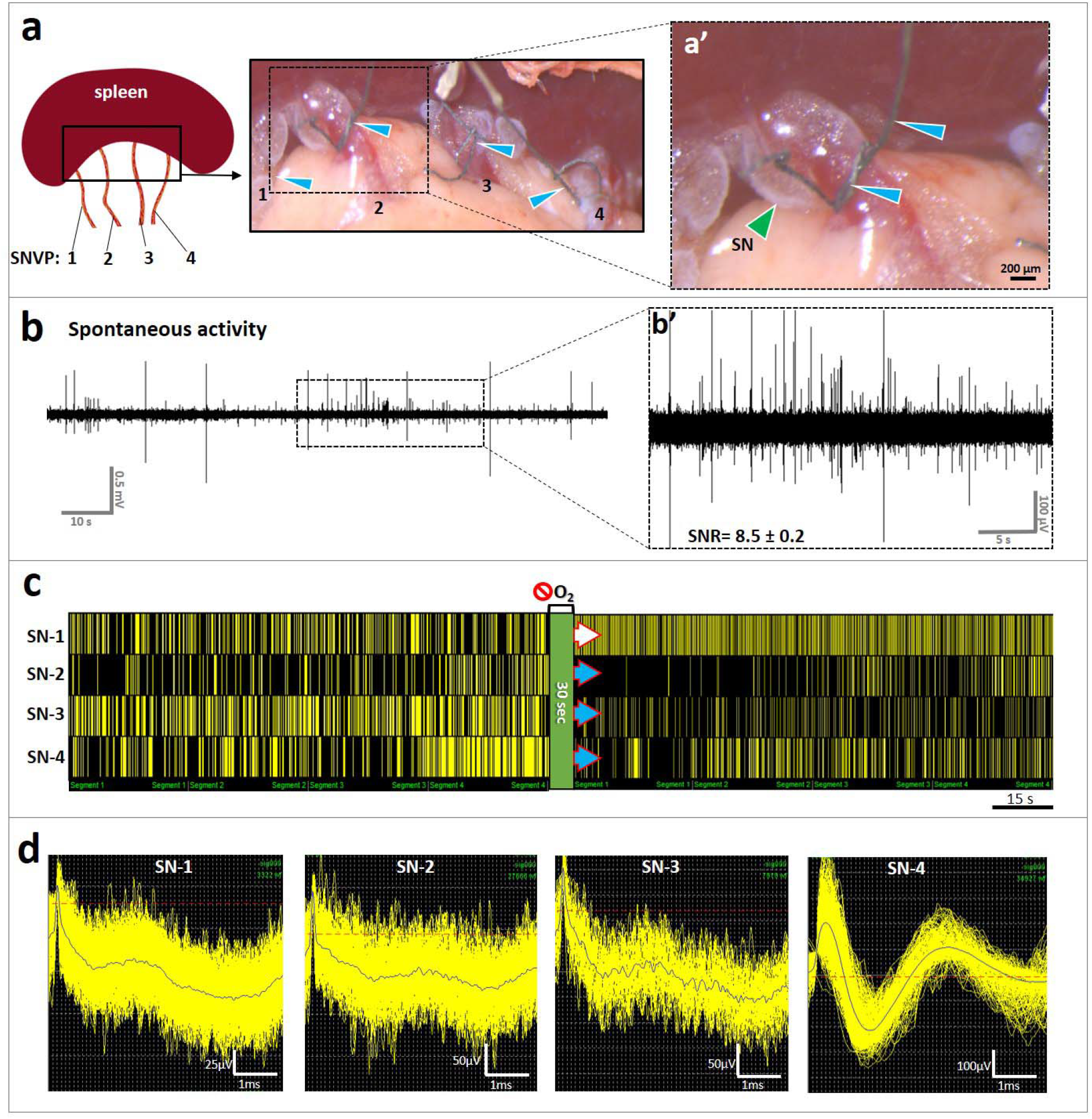
Sutrode recording of SNVP terminal branches revealed differential response to oxygen deprivation. a) Schematic and pictures of the SNVP branches (1-4; apical to basal) interfaced with sutrodes (blue arrowheads in a and a’), tied around the SNVP (a’), green arrowhead points out a SN branch. b) Raw spontaneous recording of neural activity from SN-1 (signal to noise ratio, SNR= 8.5 ± 0.2 peak to peak, b’). c) Simultaneous raster plots from the four SN branches showing differential patterns of activity before and after Oxygen deprivation (green strip) of identified CNAPs waveforms (d). The spontaneous activity increased in SN-1 (white arrow), but decreased in SN-2, 3, and 4 (blue arrows). SN, splenic nerve; SNVP, splenic neurovascular plexi.

In response to hypoxia, the activity in SN-1 increased in frequency, whereas SN-2 showed a sharp decrease in activity immediately after hypoxia (Fig. 1c). In order to confirm the differential functional response of the SN terminal nerve fascicles to physiological events, we used the sutrode to record the response to NPS-induced vasodilation and hypotension, from the cVN and the SN terminal branches, simultaneously (Fig. 5). Consistent with the differential axonal composition, we were able to discriminate multiple CNAP waveforms in the cVN and the SN terminal branches, with unique temporal resolution. Immediately after NPS administration, a tonically active waveform in the cVN (Wf-a, Fig. 5a) drastically reduced its firing activity (Fig. 5b), this event was followed by a delayed increase and decrease in Wf-b, and a late appearance of Wf-c 250 seconds later. Considering a threshold of 0.4 spikes/sec to interpret significant changes in activity frequency, the SN-1 nerve responded immediately after NPS and continue to increase in frequency over time. The activity levels of SN-2 and SN-4 showed an increase in frequency that coincided with that of Wf-b in the VN (yellow dotted line Fig. 5 b-d). In contrast, neural activity in SN-3 did not change substantially. Together, the physiological response from the individual SN branches suggests that they carry different functional information to and from the spleen, in a unique temporal pattern.

**Figure 5.**
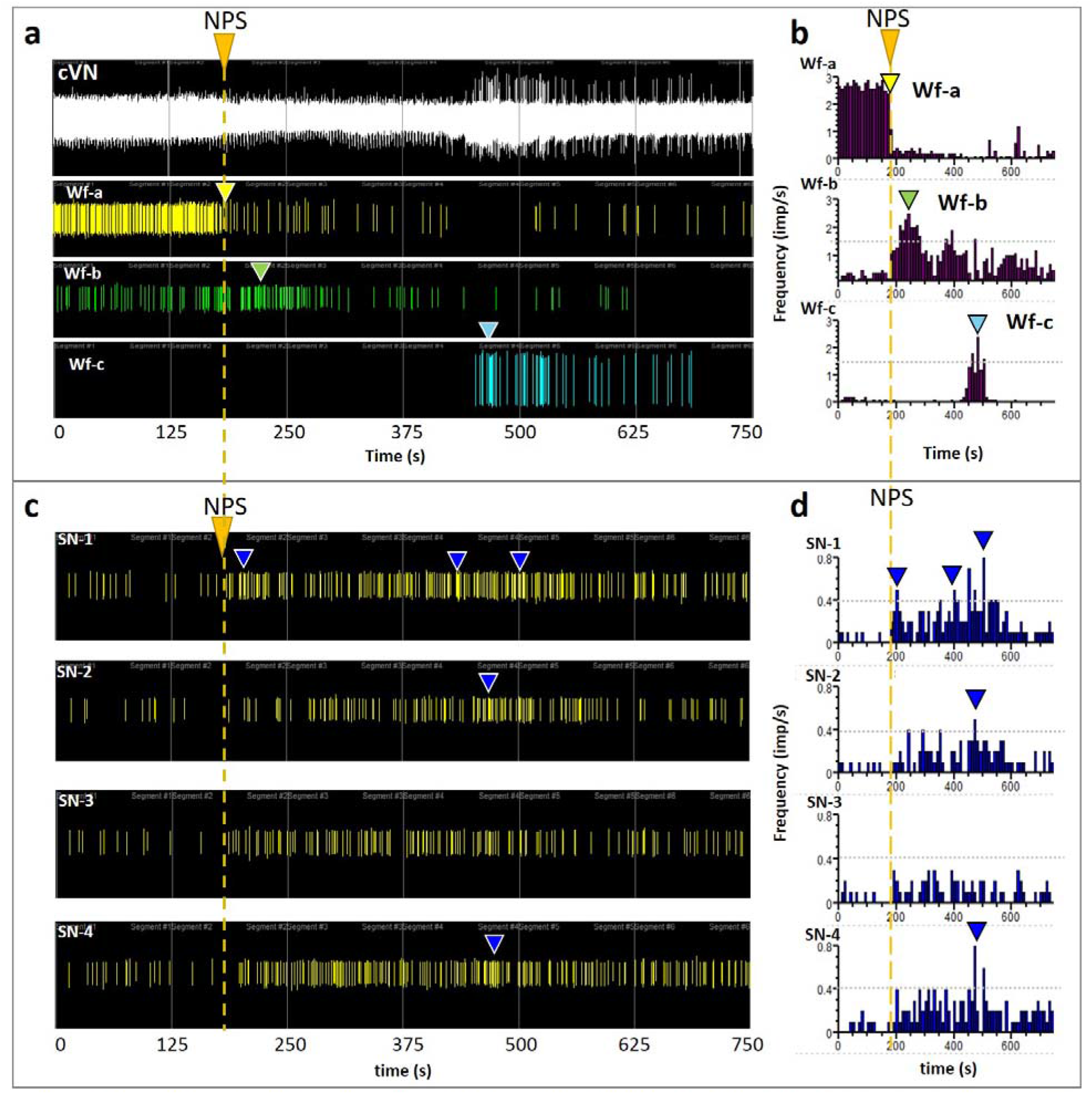
Hypotension inhibits cVN activity, followed by differential splenic branch activity. a) NPS administration (orange arrow and dashed line) causes a reduction in cVN activity (Wf-a, yellow arrow), and activation of two separate Wf (Wf-a and b, green and blue arrows, respectively), as demonstrated by the frequency histograms (b). c) Simultaneous recording of neural activity in the four splenic branches showed a differential temporal response in frequency firing, with SN-1 following the loss of cVN Wfa, and SN-2 and SN-4 activated simultaneously with VN Wf-c. d) Frequency histograms show the specific time of increased activity over a 40% threshold (blue arrowheads in c and d). NPS, nitroprusside; cVN, cervical vagus nerve; Wf, waveform; SN, splenic nerve.

### VNS evokes differential activity in SN terminal branches

The apparent temporal correlation between induced activity in the VN and the terminal SN branches indicated a direct VN control on the spleen. Such functional connection between the VN and the spleen has been suggested before, but it is currently a topic of controversy. Given the differential anatomical and functional activity in the four SN terminal branches, we reasoned that the functional connection of the VN to the spleen might be related to some, but not all axons in the SN, possibly explaining the controversial reports. To address this question, we recorded the neural activity from the four SNVP in response to cVN stimulation. The application of 30 seconds stimulation of 0.5V pulses, evoked an immediate 3-fold increase in nerve frequency and amplitude in SN-1 and some in SN-2 branches, but no changes were seen in SN-3 and SN-4 (Fig. 6a-b). This activity was blocked with lidocaine applied to the splenic branches confirming that this activity was neural in nature (Fig. 6c). The differential response was also evidence by increasing the stimulation voltage. At 1.2V the activity in the VN increases, followed by increase neural activity in SN-1 and SN-2. Activity in SN-3 only increased with a 2V stimulation and, in contrast, that of SN-4 tended to decrease (Supplementary Fig. 2).

**Figure 6.**
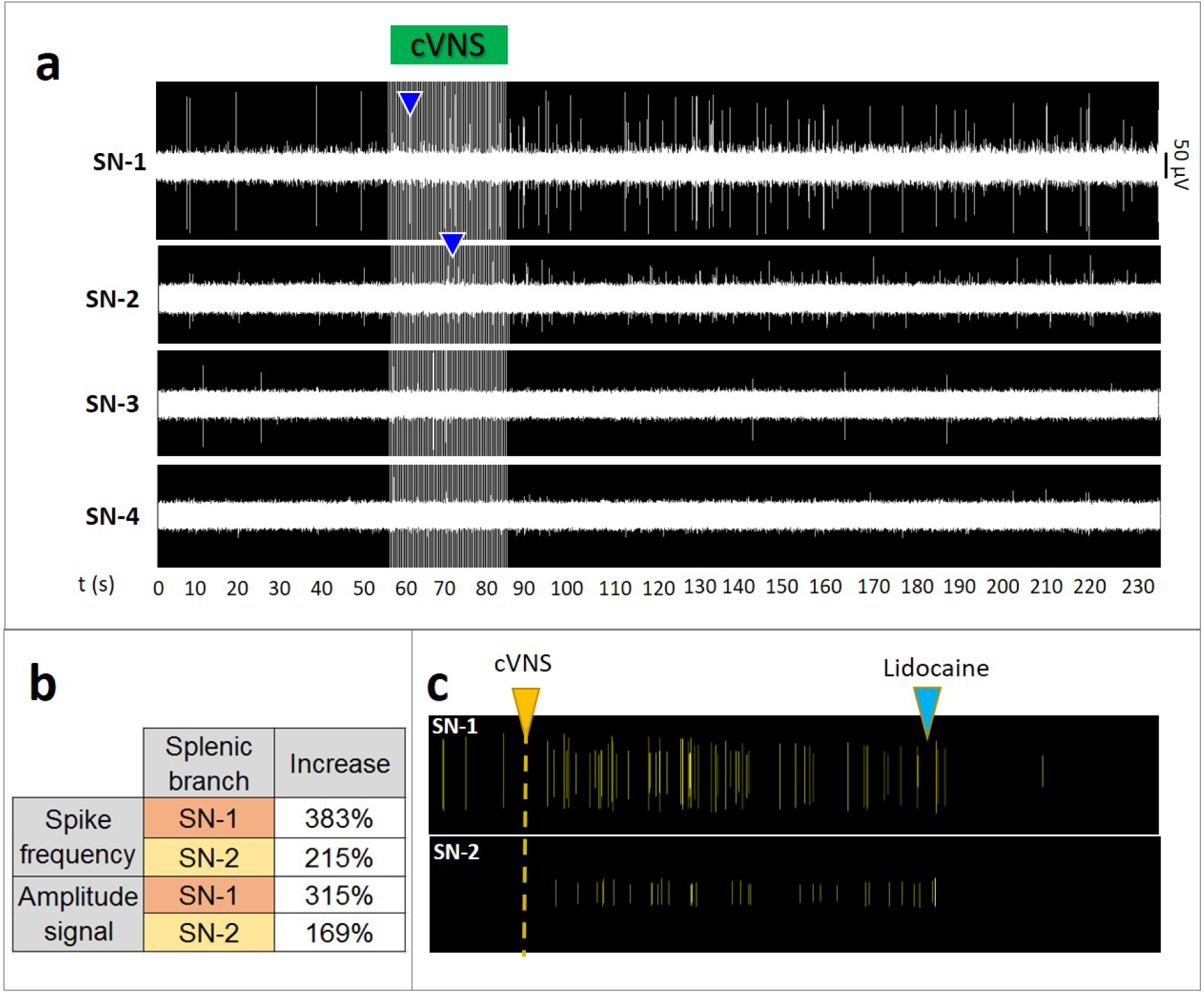
Apical-to-ventral activity gradient in SN branches in response to cVNS. a) cVNS with 0.5V pulses, 30s pulses evoked an immediate increase in SN-1 and SN-2 (blue arrowheads), and less in SN-3. No activity was evoked in SN-4. b) Spike frequency and amplitude signal increase for SN-1 and SN-2. c) lidocaine (blue arrow) blocked the neuronal activity evoked by cVNS (yellow arrow), confirming its nature. SN, splenic nerve; cVNS, cervical vagus nerve stimulation.

To better establish the functional relationship of the VN and the SN terminal branches we used sutrodes to record form the SN-1 in response to stimulation of the cVN and the subdiaphragmatic VN (SD-VN) (Fig. 7). We confirmed that cVNS evokes a response in SN-1 that increases in frequency firing (Fig. 7a). However, increasing the depolarization voltage to 1.5V did not seem to have the same effect, even if the pulse duration is increased to 0.5 ms. Importantly, SD-VN stimulation at 80mV for 0.2 ms pulses also induced increased activity in SN-1 within 10 seconds. This activation was parameter specific, since a progressive increase in stimulation voltage to 100 mV resulted in inhibition of the evoked response (Fig. 7b). These results underscore the importance of parameter specification in the control of activity of the splenic terminal plexi.

**Figure 7.**
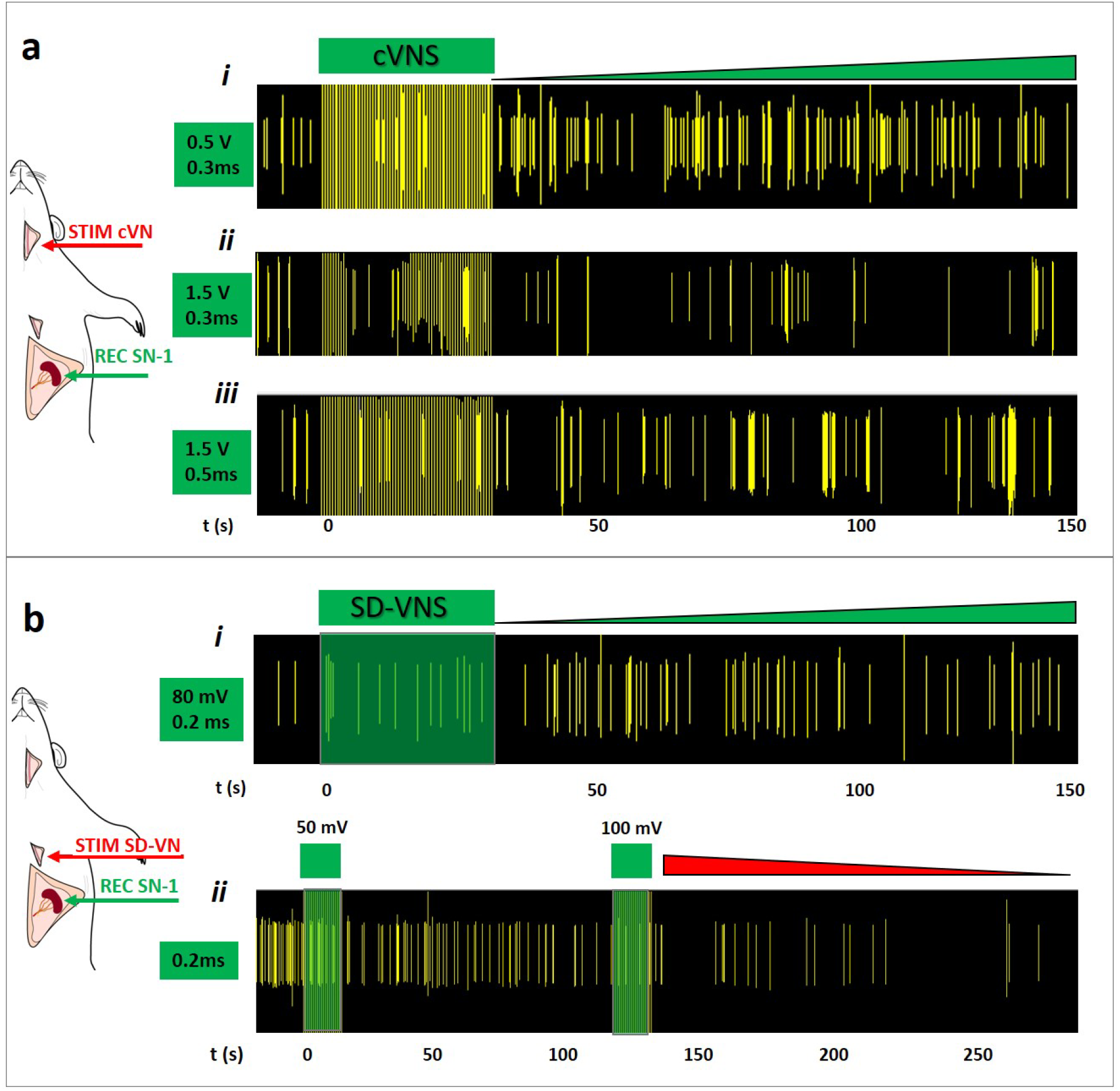
Inhibition of apical branch activity by cVNS. a) The activity in SN-1 after cVNS increased at: *i)* 0.5V. *ii)* At 1.5V cVNS caused the opposite effect. *iii)* Increasing the pulse to 0.5 ms, at 1.5V had an intermediate effect. b) SD-VNS with: *i)* 0.2 ms stimulation at 80mV caused an increase in activity, whereas stimulation of 50 and 100 mV (*ii*), inhibited SN-1 spontaneous activity. Green and red triangles indicate an increase and decrease in activity, respectively. cVNS, cervical vagus nerve stimulation; SD-VNS, subdiaphragmatic vagus nerve stimulation; SN, splenic nerve.

### Tracing of VN fibers to the splenic terminal plexi

The activity recorded in the SN-1 splenic branch, seconds after the SD-VN stimulation, suggested the possibility that this branch is directly connected to the VN. To evaluate this possibility we used viral tract-tracing by injecting an adenovirus (AdV) encoding the green fluorescent protein tracer (Ad-GFP) to the SN-1, and a second AdV tracer with the red fluorescent label mCherry to SN-3 (Ad-mCherry), and looked for labeled axons in the cVN 6 days after (Fig. 8a). The expression of the GFP biomarker was confirmed in the spleen where clusters of small axons were observed (Fig. b-c’). This tracer was also found sparsely in the cVN, where clusters of axons were also visible (Fig. 8d-d”), indicating a direct innervation of the SN-1 by the VN, albeit in very small numbers and not easily detected. A similar pattern was observed with the Ad-mCherry signal, as the expression of the fluorescent protein was confirmed in the spleen and small axons were seen in the cVN fascicles (Fig. 8e-e”). Together, the data supports the notion that VN innervates the spleen directly, albeit sparsely.

**Figure 8.**
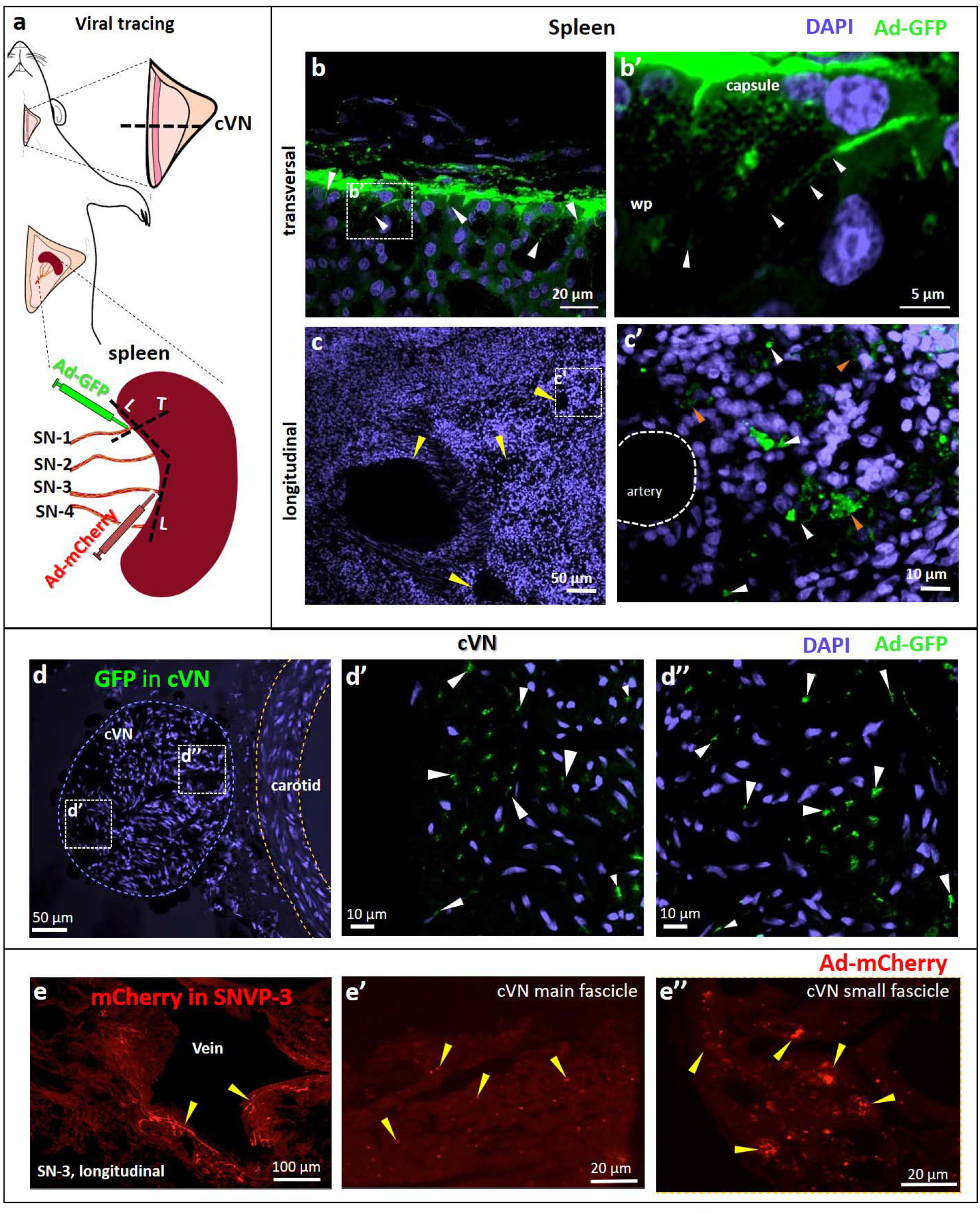
Apical and basal SN branches have direct projections to the VN. a) Schematic of Ad-GFP injections in the SN-1, and Ad-mCherry in SN-3 for tract-tracing. b) Transverse histological sections of the spleen showing GFP+ cells (white arrows) in the splenic capsule and white pulp (wp). The white square is magnified as b’. c) Transverse sections show blood vessels (yellow arrows) in proximity to GFP+ nerve fibers (white arrows in c’), often organized in clusters (orange arrows). d) Cross-section of cVN, adjacent to the carotid artery (blue and yellow dotted lines); magnifications presented as d’ and d’’ show two GFP+ axonal clusters. Cellular nuclei were stained with DAPI. e) mCherry+ axons (yellow arrowheads) adjacent to a vein in the spleen adjacent to SNVP-3; e’ and e” show mCherry+ axons in the cVN.

## Discussion

Neural regulation of the immune system is critical for an effective and measured response to infection, trauma, or injury, and deregulated responses contribute to chronic inflammatory conditions ^24^. Neuromodulation of the spleen, inhibits the production of inflammatory cytokines and has substantial clinical applications including rheumatoid arthritis, colitis, and sepsis ^25-27^. However, the neural activity in the main splenic nerve trunk is only partially understood and that of the terminal branches have remained unexplored due to technological challenges to place highly sensitive and flexible electrodes on the neurovascular plexi. Previous recordings from the main SN required the surgical isolation of the nerve from the blood vessels, and resection of the nerve before entering the spleen in order to facilitate the implant of the SN with traditional hook electrodes. Such studies reported a 27% increase in splenic vascular mechanoceptor afferent activity over 5 min, in response to an increase in splenic venous pressure ^15^. However, the simultaneous recording of nerve activity from all splenic branches has not been previously achieved. In this work, we report the use of Pt-rGO fiber electrodes^21^ as sutrodes, for the successful recording of neural activity from all SNVP. Previously, this has been extremely difficult due to the small diameter of these fascicles in rodents (i.e., 50-100 μm nerves with 300- 400 unmyelinated axons) and their location as they travel between blood vessels and are surrounded by fatty tissue. In the rat, terminal splenic fascicular branches are 8-12 times smaller and with approximately 1.75 % of the number of axons compared to the cVN (600-800 μm; >20,000 axons). The combination of a small and fragile nerve, and the low axon content, makes the detection of CNAP from splenic terminal neurovascular plexi, a very challenging task. Here we report that the unique flexibility, mechanical strength, and sensitivity of the sutrode, allowed its placement over the SN terminal plexi, and the recording of physiologically and electrically evoked CNAPs from the terminal SN fascicles.

Placing the sutrode on the cVN allowed the recording nerve activity evoked hypotension and hypoxia. This capability is not surprising as recent multi-contract cuff electrodes^28^ or intraneural carbon nanotube yarn electrodes can also be used to record CNAP from the cVN ^29^. However, the information recorded in those studies is limited and often requires the use of advanced signal processing methods for the identification of clustered waveforms ^30^. In this study, we were able to detect multiple waveforms associated with physiological changes directly. This is clearly shown in Fig. 5a where an immediate reduction in cVN activity was observed after NPS-induced vasodilation, suggesting the inhibition of afferent tonic activity likely from low-threshold vascular mechanoceptors ^31^. Thus, the use of the sutrodes increased the quality and temporal resolution of the recorded neural information.

The unique flexibility and sensitivity of the sutrode allowed for the first time the interfacing of the SNVP terminal branches and revealed that defasciculation from the main SN into the branches is not functionally homogeneous. We learned that each terminal fascicle has unique neuron fiber content and responds selectively to specific physiological events. Since the vast majority of the nerve fibers in the splenic nerve are efferent sympathetic originating in the celiac-superior mesenteric ganglia ^32^, the evoked increase in CNAP activity by hypotension and hypoxia is likely sympathetic. However, the nature of the parasympathetic content and control has been a subject of debate and this study offers a new reconciliatory perspective.

It is well known that the spleen is under reflex modulation by the VN mediated by the parasympathetic humoral arm, and afferent activity in response to ventral spleen compression ^14^. Most recently, the SN has been shown to be a necessary component of the neuroimmune reflex circuit with the vagus nerve, which is modulated by inflammatory cytokines such as TNFa, and IL-1 ^6,33,34^. However, previous denervation and tract-tracing studies using traditional retrograde fluorescent molecules, have failed to confirm a direct innervation from spleen to the VN ^35,36^, and electrophysiological studies have not confirmed a direct VNS evoked activity in the SN ^9^. The limitation of these previous studies is the use of silver wire electrodes for the SN recordings, and the lack of spatial resolution by the traditional Fast-blue tracers used in those studies^9^. In this report, the use of the highly sensitive and flexible Pt-rGO fiber electrodes allowed the recording of neural activity from intact splenic branches and showed evoked responses within 10 seconds after cVNS in some branches, and delayed reflective activity in others (Fig. 6). The discovered heterogenous fiber content and diverse temporal response to physiological or electrically evoked activity in this study, suggest that recording from previous reports was limited by the positioning and sensitivity of traditional hook electrodes on the SN, which only samples part of the neural activity. In addition, our viral tract tracing revealed a small number of axons from the spleen in the cVN, which may explain on why these have been missed in previous reports using traditional axon tract-tracing methods with limited spatial resolution and electrophysiological recording with low temporal resolution^9^. Although the combined anatomical and functional connection between the VN and the spleen was indicated by our results, further studies are warranted to confirm this direct VN-spleen pathway and determine the location and nature of the cells of origin and targeted cells.

The potential clinical impact of selective neural stimulation of the splenic nerve terminal branches has been suggested recently, as stimulation of a terminal apical branch (not associated with a blood vessel; 650 μA, 100 μs pulses at 10 Hz), was shown to increase the release of acetylcholine in the spleen and inhibit inflammatory cytokine secretion^13^. In this study, we expand and further define this possibility, by demonstrating that the type of terminal branch, and the specific electrical stimulation parameters used, can either increase or suppress the activity on the splenic fascicles, paving the way for a more selective and complete neuromodulation strategy for the spleen and, consequently, for the treatment of chronic inflammatory conditions.

Together, this study reports the use of a flexible, mechanically robust, and highly sensitive Pt-rGO sutrode for the interfacing of small neve fascicles in neurovascular plexi. Their use allows for the first time the interrogation of small nerves, despite their inter-vascular location and small size. This approach will likely expand the recording of other neurovascular plexi, particularly those in internal organs, to inform, facilitate and enable their neuromodulation for bioelectronic applications.

## Materials and Methods

### Animal use

#### Ethics statement

All protocols and surgical procedures were designed to prevent animal discomfort and suffering at any moment. These were approved by The University of Texas at Dallas, Institutional Animal Care and Use Committee (IACUC, protocol No.14-09), and follows the guidelines provided by the National Institute of Health (NIH).

### Surgical procedures

A total of 31 Female Sprague Dawley rats (300-350 g; Charles River, Wilmington, MA) were used for the experiments. The animals were anesthetized with vaporized isoflurane (2%) in a constant oxygen flux (2 L/min) delivered by a calibrated vaporizer and maintained throughout the experiment. The animal temperature was maintained with an electrical warm pad. Anesthesia and vital signs were monitored constantly with standard methods throughout the experiment. Individual nerves studies included: *-Sciatic nerve (ScN)* exposed by a 4cm longitudinal incision on the animal’s hind limb from hip to knee following the femur. The biceps and quadriceps femoris were separated to visualize the ScN. - *The cervical vagus nerve (cVN)* was exposed by making a longitudinal medial incision in the anterior part of the neck at the cervical level, using the anterior sternum as reference. The sternomastoid muscle was separated in oblique orientation to the midline and the cVN identified lateral to the carotid artery. - *Sub-diaphragmatic VN* (SD- VN) was isolated by a midline incision (2.5cm) made on the abdominal wall, the stomach was moved slightly to expose the esophagus, where the SD-VN trunks were identified between the diaphragm and the gastric cardia. - *The spleen* was identified in the upper left portion of the abdomen under the left part of the stomach. The splenic neurovascular terminal plexi were evidenced by gently lifting the spleen (Supplementary Video 3). For the ScN, cVN, and SD-VN, the connective tissue was carefully removed prior to electrode implantation. At the end of the experiments, an overdose of sodium pentobarbital (120 mg/kg) was used for euthanasia.

### Platinized graphene electrodes fabrication and validation

The rGO fiber electrodes (aka sutrodes) were fabricated by the wet spinning of GO and characterized as previously reported ^21^. Briefly, GO was reduced in hypo-phosphorous acid solution (50% in water, Sigma-Aldrich) at 80 °C for 24 h. After washing and drying, the rGO fibers (40 µm diameter) were sputter-coated with a 200 nm Pt layer. The Pt-rGO fibers were cut into 10 cm pieces by dipping into liquid nitrogen for about 1 min and then cut with a pre-cooled scissor. One end of the fiber was welded to a Pt wire, and 1 cm in the other end was covered with Parafilm, before coating with Parylene C using a deposition system (Specialty Coating System, PDS 2010 Labcoater). The Parafilm was then carefully removed to expose the Pt-rGO at the recording end of the fiber. The ultrastructure topology was assessed by scanning electron microscopy (SEM) using a microscope JEOL JSM-7500FA.

The electrical conductivity of Pt-rGO fibers was measured using a digital multimeter (Agilent 34401A) and the impedance documented while making the sutrode into a knot (Supplementary Video 1). We then measured the baseline electrical noise recorded with the sutrode in saline solution and compared it to that of a commercial stainless-steel bipolar hook electrode (PBAA15100, FHC Inc., Bowdoin, ME) (n=6).

### Electrode implantation

To implant the sutrodes, the nerves were gently lifted using a blunted glass rod. A small piece of waxed parafilm was placed underneath the nerve to isolate it from surrounding tissue. Fine angled microsurgical forceps were used to wrap the Pt-rGO fiber electrode around the target nerve (SNVP-1 to 4, ScN or cVN) and an overhand knot was carefully tied, making sure that the circulation on the epineurium microvasculature was not obstructed. To stimulate the ScN, the cVN and the SD-VN, a hook electrode (PBAA15100, FHC Inc., Bowdoin, ME) was used. Specifically for the spleen surgery, saline humidified gauzes helped to separate and maintain hydrated surrounding visceral tissue.

### Mean Arterial Pressure and Drug delivery

The mean arterial pressure (MAP) was measured with a catheter in the femoral artery, and the drug delivery was administrated in the femoral vein. n=6 rats were used for this study, 1 to 3 physiological tests per rat. A 1.0-1.5 cm incision in the medial aspect of the leg was made to expose both vessels, and a cannula (0.6mm outer diameter) previously filled with heparinized saline (20 IU/mL) was inserted and secured to the muscle using 4.0 silk sutures. The arterial cannula was connected to a previously calibrated pressure transducer (AD-Instruments, MLT1199) coupled to a bridge amplifier and power supply modulus (AD-Instruments, FE221 and ML826, respectively) for continuously MAP evaluation. The venous cannula was coupled to an infusion system to administrate the vasoactive drug, nitroprusside (5mg/mL, a bolus of 2.5 µg/g weight; 71778 Sigma Aldrich). Simultaneous nerve activity was recorded with the sutrodes placed on the cVN and SNVP-1 to 4. PowerLab data acquisition system (AD Instruments, Colorado Springs, CO) and LabChart Pro (AD Instruments, Colorado Springs, CO) softwares were used to process and analyze the mean arterial pressure data.

### Electrophysiology

Graphene electrode fibers were coupled to a customized adapter and connected to a 20X amplifier coupled to an Omniplex data acquisition system (Plexon Inc. Dallas TX). The electrophysiological activity was recorded at 40KHz. A reference electrode consisted of a graphene fiber placed between the sternomastoid muscle and the skin or under the abdominal skin tissue for the VN or SN branches respectively. Confirmation of the neural nature of the recorded activity was done by reduction of the signal after topical lidocaine application on the nerves, and elimination of the signal after euthanasia. Off-line sorter software (Plexon Inc, v3.3.5) and NeuroExplorer software (Nex Technologies, v 4.135) were used for data analysis.

#### ScN recordings

The sciatic nerve was implanted with a sutrode and used to record evoked activity by 1) applying electrical stimulation with a bipolar hook electrode, placed 3mm far in the proximal portion. 0.6V symmetrical biphasic pulses, 0.3 ms width at 2 Hz with not interphase delay were applied. n=6 rats were used for this study, 3 to 8 electrophysiological tests per rat. Or by 2) implanting 2 sutrodes in the biceps muscle (∼20 mm distanced from the recording site), using them as conductive sutures, and applying symmetrical biphasic pulses, 0.3 ms width at 2 Hz with not interphase delay, which elicited tissue contraction (n=6 stimulation tests in one animal).

#### VN Stimulation

Evoked activity in the SN-1 to 4 was evaluated by the stimulation of the cVN. n=3 rats were used for this study, 3 to 5 tests per rat were performed with a total of 11 evoked responses. SN-1 to 4 were implanted with sutrodes and the cVN with a hook electrode for bipolar stimulation. 0.3 ms pulses of 0.5V symmetrical biphasic pulses, 1ms inter-pulse delay were applied by 30s at 2 Hz. Furthermore, the study was focused on SN-1 activity. cVN was stimulated by 30s in different experimental sets: i) 0.3 ms pulses, 0.5V; ii) 0.3 ms pulses, 1.5 V, and iii) 0.5 ms pulses, 1.5 V. Finally increased electrical trains of stimulation were applied to the cVN with a hook electrode, in this experiment a sutrode was included in the cVN, to simultaneously with the SN-1 to 4, record the evoked activity. 0.3 ms biphasic pulses, at 2 Hz, during 30 s were applied: 1.2, 1.4, 1.6, 1.8, and 2.0V. 3.5 min were allowed between stimulation sets.

SD-VN stimulation effect on the SN-1 implanted with a sutrode was evaluated. n=4 rats were used for this study, 3 to 4 experiments per rat were performed (total SD-VN stimulations=13). Different stimulation parameters were applied with a hook electrode: 0.2 ms pulses, 2 Hz, at 50, 80 and 100 mV.

#### Oxygen reduction

Evoked neural activity in response to Oxygen restriction was evaluated by implanting sutrode recording electrodes on SNVP-1 to 4 or cVN. n=5 rats were used for this study, 1 to 5 experiments per rat were performed (total physiological tests=17). Baseline was recorded, followed by 2 min of Oxygen deprivation, elicited by closing the supply source. A hermetic face mask and extra latex seal in the borders between the mask and the head of the rat were used for this experiment in an effort to secure the Oxygen reduction. After opening the Oxygen flux, continuous recordings were collected. Our Oxygen deprivation system was validated by using a calibrated fiber-optic oxygen micro-sensor (OxyLite, Oxford-Optronix), which in saline bubbling Oxygen from the system used for the *in-vivo* experiments, allowed to determine that ∼36 s were required for the Oxygen to drop after closing the Oxygen source.

### Ultrastructural analysis

The ultrastructural analysis of the main splenic nerve and the neuro vascular plexi of the spleen was assessed by transmission electronic microscopy (TEM). Four rats were trans-cardialy perfused with physiological saline solution (NaCl, 0.9%), followed by a fixative solution that consists of 4 % paraformaldehyde, 0.5 % glutaraldehyde, in sodium arsenate (cacodylate) buffer (0.1 M, pH 7.4). The nerve sections of interest were attached to silk sutures for easy manipulation and dissected for post-fixation. The samples were post-fixed in 3.0 % glutaraldehyde in cacodylate buffer (0.1 M pH 7.4), at 4 °C and processed for resin embedding and ultrathin sections. Standard toluidine blue staining was performed on 700 nm semi-thin sections to evaluate the structural organization. Ultra-thin sections of 60-70 nm were obtained using an ultramicrotome, mounted on copper grids, and coated with uranil acetate and silver citrate for contrast. The sections were imaged using a transmission electron microscope (TEM, JEOL 1400 Plus, JEOL, USA). The images were obtained at high magnification and axon diameters were measured using the Image J software (NIH).

### Adenoviral neuronal tracing

Two recombinant serotype 5 adenovirus (AdV) that codify for the expression of mCherry (Ad-mCherry) and GFP (Ad-GFP) proteins, under the control of the CMV promoter (Vector BioLabs, Philadelphia, USA) were used as axonal tracers. These were administered (□3.0□ ×□ 10^7^ viral particles/μl) with a Hamilton Neuro syringe (Hamilton, 1710) into the splenic terminal branches. Ad-GFP was used on SN-1 and Ad-mCherry on SN-3 (n=2). Extreme efforts were made to prevent the spread of the tracers to the surrounding tissue and all the procedures were made using a surgical stereoscope. A parafilm was placed underneath the neurovascular-fat tissue during the administration, 15 min were allowed for the virus to be incorporated into the tissue after the administration, then a mini cotton-tip was used to remove the viral solution and physiological interstitial liquid. Finally, a microneedle was used to make 5 washes with saline to maximize the removal of viral particles not incorporated in the tissue; each wash was followed by cleaning with a cotton tip. The parafilm was removed and the incision closed, muscle with 4.0 silk sutures, and the skin with metallic staples. Cephazolin (5mg/kg, IM) and buprenorphine (1mg/kg, SQ) were administrated subsequently as antibiotic and analgesic, respectively. After 6 days, the rats were intracardially perfused with physiological saline solution (NaCl, 0.9%), followed by fixative solution (4 % paraformaldehyde in PBS). The VN attached to the carotid artery, and the SN branches were isolated and post-fixed in the same solution for 2 h. The samples were cryoprotected in increased sucrose gradients (10, 20, and 30%), embedded in OCT compound (Sekura Finetek, Torrance, CA, USA), and sectioned (45 μm thick) in a cryostat (Leica, CM3050s). The cellular nuclei were contrasted with 4′,6-diamidino-2-phenylindole (DAPI, 1Lmg/mL) and imaged in a confocal microscope (Nikon Eclipse Ti, Japan).

### Statistics

Kolmogorov-Smirnov normality test was used to determine the difference in axon diameter population in the different groups, followed by linear and non-linear regression of the normalized axon diameter distribution, and used the slope m to determine the statistical significance among the different splenic nerve terminal branches. Statistical analysis was run on Prism 8 software (GraphPad, San Diego, CA).

## Acknowledgements

The authors would like to thank the Australian National Fabrication Facility (ANFF) Materials node, for access to services and equipment. We acknowledge Prof. Rouhollah Ali Jalili for contributions in the standardization and fabrication quality of the Pt-rGO fiber electrodes. Prof Gordon Wallace and Kezhong Wang gratefully acknowledge funding from the Australian Research Council Centre of Excellence Scheme (Project Number CE 140100012). This study was supported by intramural funding at UT Dallas.

## Author contributions

Design of the research: MRO, MAGG. *in-vivo* studies and electrophysiology: MRO, MAGG. TEM imaging: GB. Fabrication and *in-vitro* characterization of Pt-rGO: GW, KW. Analyzed the results and reviewed the manuscript: MRO, MAGG, GSB, KW, GW. Wrote the paper: MRO, MAGG.

## Conflict of Interest

MRO owns shares in RBI Medical, a medical device company. RBI Medical did not have any in role in data collection, analysis, or the manuscript. The remaining authors have no conflicts of interest.

## Supplementary Information

**Supplementary Figure 1.**
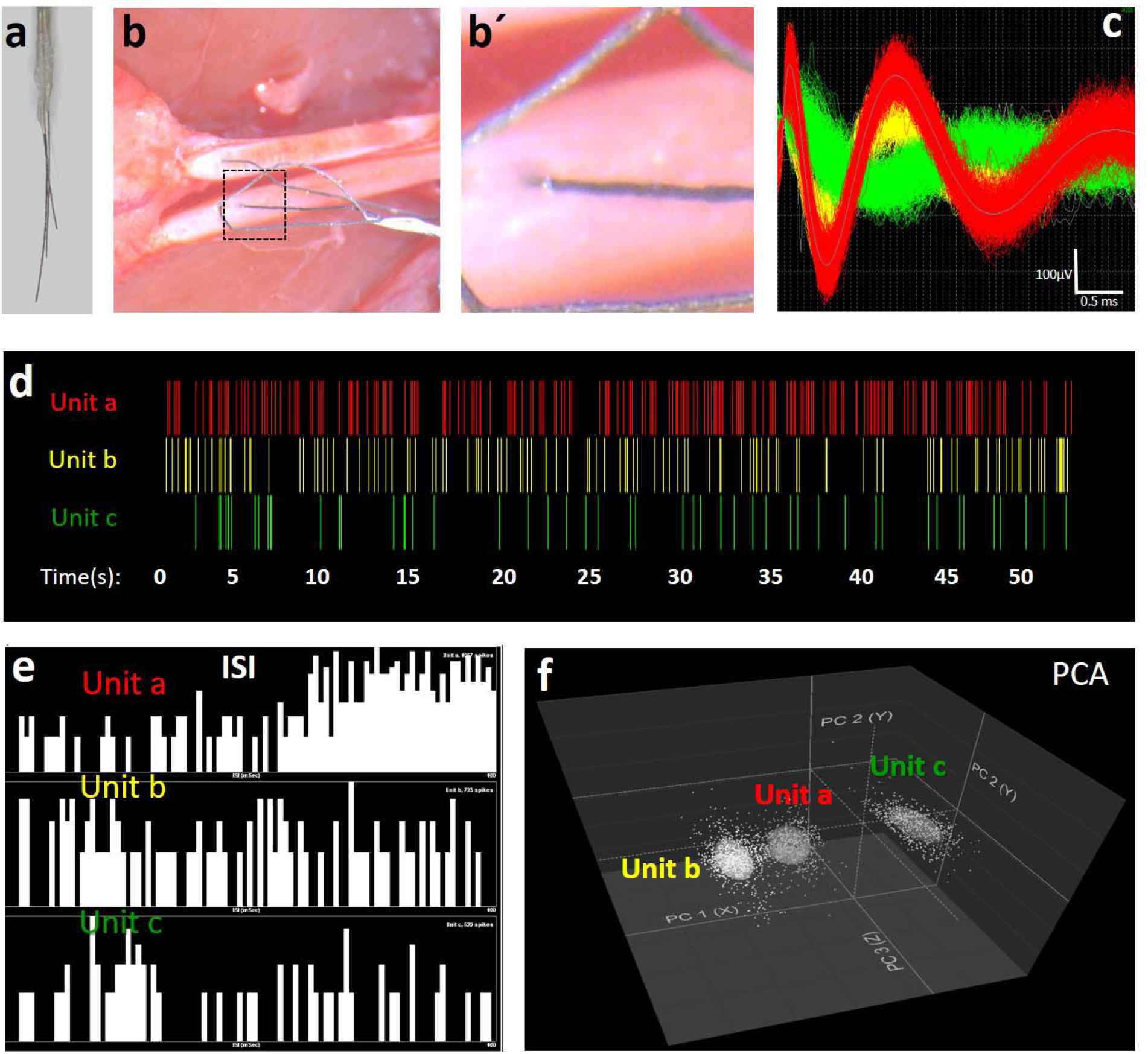
Sutrode multi-array as intraneural peripheral interface. a) Sutrode multi-array; b) sutrode multi-array implanted intraneural in two fascicles of the ScN, magnification is displayed in b’. c) representative recording from one of the electrodes identifies three single units. Raster plots, inter-spike interval analisis (ISI) and principal component analysis (PCA) are presented in d-f, respectively. SNR was calculate as 9.6.

**Supplementary Figure 2.**
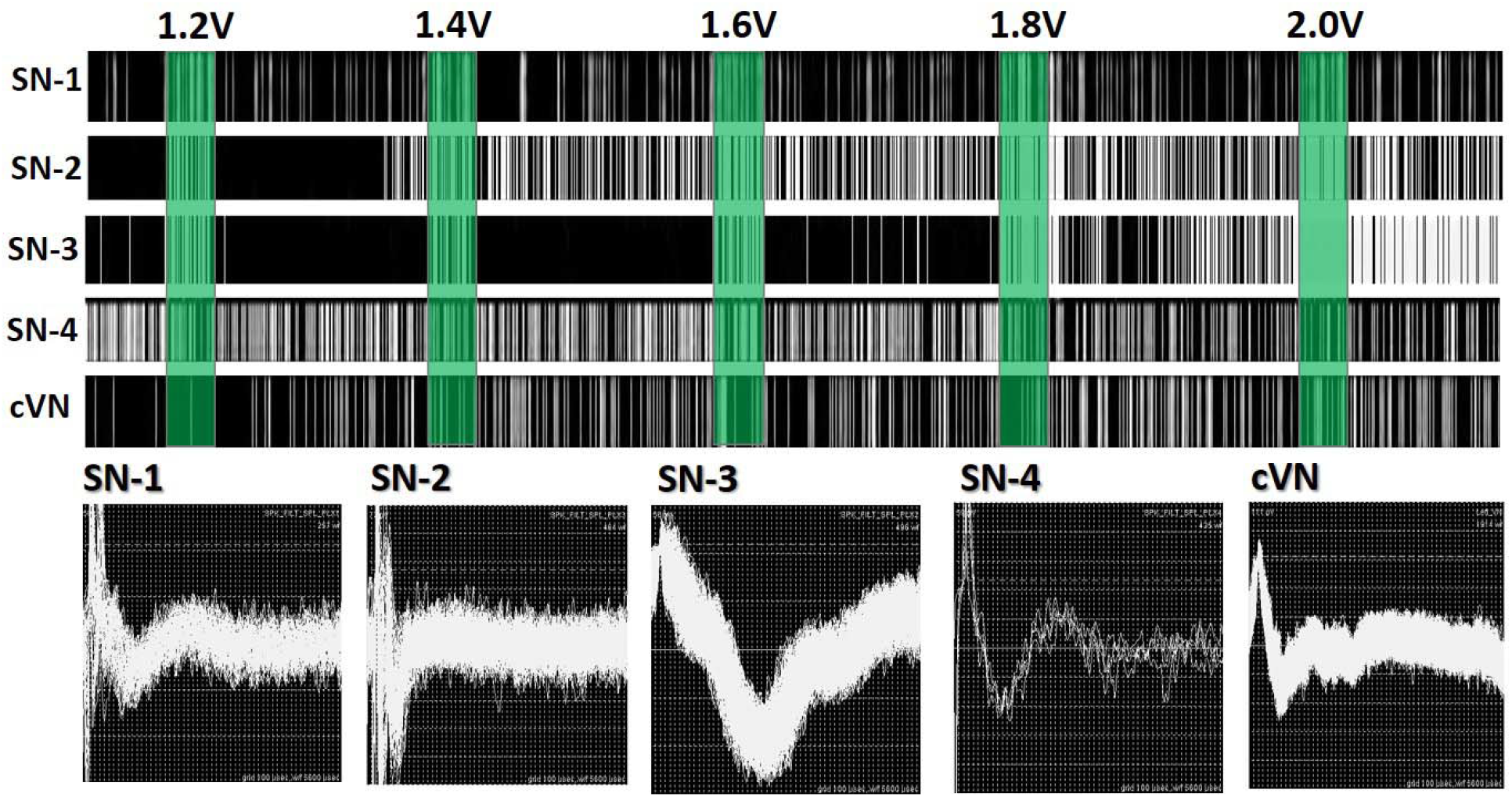
cVN stimulation evokes voltaje-dependent activity in SN terminal branches. Neural activity in SN-1 to 4 and cVN, evoked by cVN stimulation: 0.3 ms biphasic pulses, at 2 Hz, durig 30 s at 1.2, 1.4, 1.6, 1.8 and 2.0V

**Supplementary Video 1**. Sutrode flexibility and impedance stability when making a knot.

**Supplementary Video 2**. Sutrode used as a conductive suture for muscle stimulation.

**Supplementary Video 3**. Splenic neurovascular plexi terminal branches.

## Notes

### Competing Interest Statement

MRO owns shares in RBI Medical, a medical device company. RBI Medical did not have any role in data collection, analysis, or the manuscript. The remaining authors have no conflicts of interest.

## References

1 Bernik, T. R. et al. Pharmacological Stimulation of the Cholinergic Antiinflammatory Pathway. 195, 781–788, doi:10.1084/jem.20011714 (2002).

2 Borovikova, L. V. et al. Vagus nerve stimulation attenuates the systemic inflammatory response to endotoxin. Nature 405, 458–462, doi:10.1038/35013070 (2000).

3 Tsaava, T. et al. Specific vagus nerve stimulation parameters alter serum cytokine levels in the absence of inflammation. Bioelectronic Medicine 6, doi:10.1186/s42234-020-00042-8 (2020).

4 Ackerman, K. D., Felten, S. Y., Bellinger, D. L. & Felten, D. L. Noradrenergic sympathetic innervation of the spleen: III. Development of innervation in the rat spleen. Journal of neuroscience research 18, 49-54, 123-125, doi:10.1002/jnr.490180109 (1987).

5 Berthoud, H.-R. & Powley, T. L. Characterization of vagal innervation to the rat celiac, suprarenal and mesenteric ganglia. Journal of the autonomic nervous system 42, 153–169, doi:10.1016/0165-1838(93)90046-w (1993).

6 Tracey, K. J. The inflammatory reflex. Nature 420, 853–859, doi:10.1038/nature01321 (2002).

7 Rosas-Ballina, M. et al. Acetylcholine-Synthesizing T Cells Relay Neural Signals in a Vagus Nerve Circuit. Science 334, 98–101, doi:10.1126/science.1209985 (2011).

8 Koopman, F. A. et al. Vagus nerve stimulation inhibits cytokine production and attenuates disease severity in rheumatoid arthritis. Proceedings of the National Academy of Sciences of the United States of America 113, 8284–8289, doi:10.1073/pnas.1605635113 (2016).

9 Bratton, B. O. et al. Neural regulation of inflammation: no neural connection from the vagus to splenic sympathetic neurons. Experimental Physiology 97, 1180–1185, doi:10.1113/expphysiol.2011.061531 (2012).

10 Nance, D. M. & Sanders, V. M. Autonomic innervation and regulation of the immune system (1987–2007). Brain, behavior, and immunity 21, 736–745, doi:10.1016/j.bbi.2007.03.008 (2007).

11 Liu, D. L. et al. Anatomy of vasculature of 850 spleen specimens and its application in partial splenectomy. Surgery 119, 27–33, doi:10.1016/s0039-6060(96)80209-1 (1996).

12 Lorton, D. Substance P innervation of spleen in rats: Nerve fibers associate with lymphocytes and macrophages in specific compartments of the spleen. 5, 29–40, doi:10.1016/0889-1591(91)90005-u (1991).

13 Guyot, M. et al. Apical splenic nerve electrical stimulation discloses an anti-inflammatory pathway relying on adrenergic and nicotinic receptors in myeloid cells. Brain, behavior, and immunity 80, 238–246, doi:10.1016/j.bbi.2019.03.015 (2019).

14 Herman, N. L., Kostreva, D. R. & Kampine, J. P. Splenic afferents and some of their reflex responses. The American journal of physiology 242, R247–254, doi:10.1152/ajpregu.1982.242.3.R247 (1982).

15 Moncrief, K. & Kaufman, S. Splenic baroreceptors control splenic afferent nerve activity. American Journal of Physiology-Regulatory, Integrative and Comparative Physiology 290, R352–R356, doi:10.1152/ajpregu.00489.2005 (2006).

16 Gautron, L. et al. Neuronal and nonneuronal cholinergic structures in the mouse gastrointestinal tract and spleen. Journal of Comparative Neurology 521, 3741–3767, doi:10.1002/cne.23376 (2013).

17 Bellinger, D. L. et al. Neuropeptide innervation of lymphoid organs. Ann N Y Acad Sci 594, 17–33, doi:10.1111/j.1749-6632.1990.tb40464.x (1990).

18 Lundberg, J. M., Anggard, A., Pernow, J. & Hokfelt, T. Neuropeptide Y-, substance P- and VIP-immunoreactive nerves in cat spleen in relation to autonomic vascular and volume control. Cell and tissue research 239, 9–18, doi:10.1007/bf00214896 (1985).

19 Verlinden, T. J. M. et al. Innervation of the human spleen: A complete hilum-embedding approach. Brain, behavior, and immunity 77, 92–100, doi:10.1016/j.bbi.2018.12.009 (2019).

20 Silverman, H. A. et al. Standardization of methods to record Vagus nerve activity in mice. Bioelectronic Medicine 4, doi:10.1186/s42234-018-0002-y (2018).

21 Wang, K. et al. High-Performance Graphene-Fiber-Based Neural Recording Microelectrodes. Advanced materials 31, e1805867, doi:10.1002/adma.201805867 (2019).

22 Tian, H. C. et al. Graphene oxide doped conducting polymer nanocomposite film for electrode-tissue interface. Biomaterials 35, 2120–2129, doi:10.1016/j.biomaterials.2013.11.058 (2014).

23 Won, S. M., Song, E., Reeder, J. T. & Rogers, J. A. Emerging Modalities and Implantable Technologies for Neuromodulation. Cell 181, 115–135, doi:10.1016/j.cell.2020.02.054 (2020).

24 Pavlov, V. A. & Tracey, K. J. Neural regulation of immunity: molecular mechanisms and clinical translation. Nature neuroscience 20, 156–166, doi:10.1038/nn.4477 (2017).

25 Vida, G., Peña, G., Deitch, E. A. & Ulloa, L. α7-Cholinergic Receptor Mediates Vagal Induction of Splenic Norepinephrine. The Journal of Immunology 186, 4340–4346, doi:10.4049/jimmunol.1003722 (2011).

26 Ji, H. et al. Central cholinergic activation of a vagus nerve-to-spleen circuit alleviates experimental colitis. 7, 335–347, doi:10.1038/mi.2013.52 (2014).

27 Koopman, F. A. et al. Vagus nerve stimulation inhibits cytokine production and attenuates disease severity in rheumatoid arthritis. Proceedings of the National Academy of Sciences 113, 8284–8289, doi:10.1073/pnas.1605635113 (2016).

28 Plachta, D. T. et al. Detection of baroreceptor activity in rat vagal nerve recording using a multi-channel cuff-electrode and real-time coherent averaging. Conf Proc IEEE Eng Med Biol Soc 2012, 3416–3419, doi:10.1109/EMBC.2012.6346699 (2012).

29 McCallum, G. A. et al. Chronic interfacing with the autonomic nervous system using carbon nanotube (CNT) yarn electrodes. Scientific reports 7, doi:10.1038/s41598-017-10639-w (2017).

30 Zanos, T. P. et al. Identification of cytokine-specific sensory neural signals by decoding murine vagus nerve activity. Proceedings of the National Academy of Sciences of the United States of America 115, E4843–E4852, doi:10.1073/pnas.1719083115 (2018).

31 Keef, K. D. & Kreulen, D. L. Venous mechanoreceptor input to neurones in the inferior mesenteric ganglion of the guinea-pig. 377, 49–59, doi:10.1113/jphysiol.1986.sp016176 (1986).

32 Bellinger, D. L., Felten, S. Y., Lorton, D. & Felten, D. L. Origin of noradrenergic innervation of the spleen in rats. Brain, behavior, and immunity 3, 291–311, doi:10.1016/0889-1591(89)90029-9 (1989).

33 Rosas-Ballina, M. et al. Splenic nerve is required for cholinergic antiinflammatory pathway control of TNF in endotoxemia. Proceedings of the National Academy of Sciences 105, 11008–11013, doi:10.1073/pnas.0803237105 (2008).

34 Andersson, U. & Tracey, K. J. Reflex principles of immunological homeostasis. Annual review of immunology 30, 313–335, doi:10.1146/annurev-immunol-020711-075015 (2012).

35 Bellinger, D. L., Lorton, D., Hamill, R. W., Felten, S. Y. & Felten, D. L. Acetylcholinesterase staining and choline acetyltransferase activity in the young adult rat spleen: lack of evidence for cholinergic innervation. Brain, behavior, and immunity 7, 191–204, doi:10.1006/brbi.1993.1021 (1993).

36 Nance, D. M. & Burns, J. Innervation of the spleen in the rat: Evidence for absence of afferent innervation. 3, 281–290, doi:10.1016/0889-1591(89)90028-7 (1989).

